# Neonatal gut and respiratory microbiota: coordinated development through time and space

**DOI:** 10.1101/247122

**Authors:** Alex Grier, Andrew McDavid, Bokai Wang, Xing Qiu, James Java, Sanjukta Bandyopadhyay, Hongmei Yang, Jeanne Holden-Wiltse, Haeja A. Kessler, Ann L. Gill, Heidie Huyck, Ann R. Falsey, David J. Topham, Kristin M. Scheible, Mary T. Caserta, Gloria S. Pryhuber, Steven R. Gill

## Abstract

**Background:** Postnatal development of the microbiota in early life influences immunity, metabolism, neurodevelopment and long-term infant health. Microbiome development occurs at multiple body sites, each with distinct community compositions and functions. Associations between microbiota at multiple sites represent an unexplored influence on the infant microbiome. Here, we examined co-occurrence patterns of gut and respiratory microbiota in pre- and full-term infants over the first year of life, a period critical to neonatal development and risk of respiratory diseases.

**Results:** Gut and respiratory microbiota collected as longitudinal rectal, throat and nasal samples from 38 pre-term and 44 full-term infants were first clustered into community state types (CSTs) on the basis of their composition. Multiple methods were used to relate the occurrence of CSTs to several measures of infant maturity, including gestational age (GA) at birth, week of life (WOL), and post menstrual age (PMA: equal to GA plus WOL). Manifestation of CSTs followed one of three patterns with respect to infant maturity. First, *chronological*: independent of infant maturity (GA) at birth, and strongly associated with post-natal age (WOL). Second, *idiosyncratic*: primarily dependent on maturity (GA) at birth, with persistent differences in CST occurrence between pre- and full-term infants through the first year of life. Third, *convergent*: CSTs appear earlier in infants with greater maturity (GA) at birth, but after a sufficient post-natal interval their occurrence in pre-term infants reaches parity with full-term infants. The composition of CSTs was highly dissimilar between different body sites, but the CST of any one body site was highly predictive of the CSTs at other body sites. There were significant associations between the abundance of individual taxa at each body site and the CSTs of the other body sites, which persisted after stringent control for the non-linear effects of infant maturity. Significant canonical correlations exist between the microbiota composition at each pair of body sites, with the strongest correlations between more proximal locations.

**Conclusion:** Cross-body site associations of developing infant microbiota suggest the importance of research and clinical practices that focus on dynamic interactions between multiple microbial communities to elucidate and promote systemic microbiota development.

## BACKGROUND

Human life is dependent on a diverse community of symbiotic microbiota that have co-evolved with their human hosts to modulate crucial aspects of normal physiology, metabolism, immunity and neurologic function [1]. While our relationships with microbes may begin in utero, the limited microbial communities observed immediately after birth give way to densely colonized, diverse bacterial ecosystems within weeks, with early interactions between members of the microbial community and between the microbes and their human host responsible for features of postnatal development that influence future health [2-5]. The newborn infant microbiota is highly dynamic and undergoes rapid changes in composition through the first years of life towards a stable adult-like structure with distinct microbial communities of unique composition and functions at specific body sites [5-10]. Relatively little has been reported about longitudinal microbiota development or compositional differentiation across multiple body sites during this period. This is particularly true for high-risk pre-term infants, who because of immature mucosal and skin barriers, as well as underdeveloped immunity and suboptimal nutrition, are at increased risk for invasive infection and dysregulated inflammation of critical systems, namely the respiratory and gastrointestinal tracts. Serious perinatal complications in these pre-term infants result in prolonged hospitalization, treatment with antibiotics and delays in enteral feeding that influence interactions with microbes and inhibit microbial colonization characteristic of full-term infants [11].

While numerous microbial communities within individual body sites have been described [3, 9, 11-14], associations between the microbiota across multiple body sites or systems are less well studied [11, 15, 16]. A better understanding of the microbiota landscape and interactions across multiple body sites is needed to assess the influence of perturbations in one system on the microbiota of other systems. Elucidating the direct and indirect interactions of microbiota across multiple body sites presents a formidable analytical challenge. Available statistical methods vary widely in sensitivity and precision, with no consensus on the best approach [17]. Community profile data from 16S rRNA amplicon surveys is compositional, high-dimensional and generally observational, with ecological interactions between microbes often inferred rather than observed. Limited validation of these interactions through independent experiments or modeling leaves researchers without the data needed to reconstruct authentic interaction networks and to make meaningful biological conclusions [18]. The limited body of literature reporting on cross body site interactions is a testament to these challenges [11, 15, 16]. Our study leverages dimension reduction and longitudinal modeling techniques, allowing for the effects of within body site temporal development and cross-body site associations during early life to be distinguished and quantified for the first time. Furthermore, unlike previous studies that sampled the microbiome parsimoniously across body sites, our study sampled multiple body sites from a large cohort of pre- and full-term infants at frequent and regular intervals throughout their first year of life, within the crucial window of time when the microbial community maximally influences immune development and potential long-term health outcomes, including atopy, inflammatory bowel diseases and subtleties of neurodevelopment [19-22].

Here we describe and compare patterns of development of the microbiota of the nose, throat, and gut over the first year of life in 82 pre- and full-term infants (Table 1). Within the three body sites, we characterized development as a pattern of progression through microbiota community state types (CSTs), each differentiated by the abundance of specific taxa. We compared full- and pre-term infants on the basis of their progression through these CSTs and assessed the associations between the manifestation of CSTs, gestational age (GA) at birth, post-natal age as measured by week of life (WOL) and developmental age as indicated by post-menstrual age (PMA: equal to GA plus WOL). Three patterns of CST manifestation were identified. First, a chronological pattern in which CST occurrence was independent of GA at birth but a function of WOL. Second, an idiosyncratic pattern, with CST occurrence frequency primarily dependent on GA at birth. Third, a convergent pattern whereby lower GA at birth imposed a delay in the manifestation of a CST, with CST occurrence frequency in pre-term infants reaching parity with full-term infants after a post-natal interval proportional to prematurity. We demonstrate that although community composition is dissimilar between distal body sites, the abundance of various taxa and the occurrence patterns of CSTs is highly correlated across body sites. These associations cannot be entirely accounted for by the common influence of developmental or post-natal age on all body sites or by the direct transmission of bacteria between body sites, which suggest the existence of relationships between infant development and the microbiota across body sites that have yet to be defined.

**Table 1.** Demographics and Clinical Variables

| **Variables (N = 82)** | **Pre-Term**<br>**(Mean ± SD, N)** | **Full-Term**<br>**(Mean ± SD, N)** |
| --- | --- | --- |
| Gestational Age at Birth (weeks) | 29.56 ± 5.38 | 39.61 ± 4.65 |
| Gestational Age at Birth |  |  |
| 23-25 wks | 8 | - |
| 26-27 wks | 6 | - |
| 28-29 wks | 5 | - |
| 30-31 wks | 9 | - |
| 32-33 wks | 6 | - |
| 34-35 wks | 4 | - |
| Full term | - | 44 |
| Birth Weight (kg) | 1.38 ± 1.09 | 3.53 ± 1.1 |
| Sex (Male/Female) | 21 / 17 | 27 / 17 |
| Race (Caucasian/AA/Other*) | 22 / 10 / 6 | 30 / 5 / 9 |
| Ethnicity (Hispanic or Latino Y/N/Unknown) | 4 / 34 / 0 | 8 / 32 / 4 |
| Delivery method (C-section/Vaginal) | 24 / 14 | 21 / 23 |
| Hospital samples, Collected (Analyzed) |  |  |
| Rectal | 294 (252) | 43 (27) |
| Nasal | 288 (243) | 44 (19) |
| Throat | 174 (159) | 26 (13) |
| Post-discharge samples, Collected (Analyzed) |  |  |
| Rectal | 346 (331) | 479 (469) |
| Nasal | 350 (318) | 483 (433) |
| Throat | 202 (184) | 206 (182) |
| Total acute respiratory visits** | 51 | 51 |
| Number of infants had acute respiratory visit** | 20 | 24 |

\*Race group 'Other' includes those not reported race.
\*\* Samples collected during monthly visits where evidence of acute respiratory infection was observed were excluded in this analysis.

Overall, our results illustrate fundamental interactions between the gut and respiratory microbiomes in pre-term and full-term infants. Thus, this study will inform establishment of clinical criteria for potential therapeutic approaches that promote acquisition and maturation of a homeostatic global infant microbiome and mitigation of dysbiotic microbiota perturbations.

## RESULTS

### Overview of infant cohort

To characterize development of the neonatal gut and respiratory tract microbiota, we collected rectal, nasal, and throat swabs from 82 pre- and full-term infants over the first year of life (Table 1). From the 38 pre-term infants, weekly samples were collected while hospitalized in the neonatal intensive care unit from birth until discharge, and monthly samples were collected from discharge through one year of gestationally corrected age. From the 44 full-term infants in the cohort, monthly samples were collected through the first year of life, starting at birth. Samples collected during monthly visits in which evidence of acute respiratory infection was observed were excluded from this analysis. Microbiota from 1,079 gut, 1,013 nasal and 538 throat samples were characterized by 16S rRNA amplicon sequencing. At the onset, it was unclear how much additional information would be gained from interrogating both the nasal and throat sites, as opposed to the nasal site only. Accordingly, we sequenced and analyzed throat samples from an unbiased random subset of 40 subjects distributed evenly between pre- and full-term infants from the larger cohort, retaining the samples from the remaining 42 subjects for future work. As described below, the variability of the nasal and throat microbiota is significant, suggesting that additional analysis of both sites will provide unique insights into gut-respiratory interactions.

### Microbiome community state types (CSTs) summarize transient states, and stable equilibria

The microbiota community composition of the rectal, nasal and throat samples was quantified by 16S rRNA amplicon sequencing. To synthesize within-site sources of variation, samples from each body site were independently clustered into community state types (CSTs) using Dirichlet Multinomial mixture (DMM) models [23]. The DMM model sought to explain the operational taxonomic unit (OTU) compositional vector as a sample from a mixture of different canonical Dirichlet components. For each sample, the DMM model posterior probabilities indicated which Dirichlet component the observed vector of OTU counts most likely represented. On the basis of these probabilities, samples were assigned to clusters corresponding to CSTs which collapse the variation in microbiota composition into commonly observed archetypal states that serve as summary representations of the microbiota composition at each site. A robust resampling procedure (see Methods) identified 6 CSTs within the gut, 7 within the nose, and 6 within the throat, with each CST distinguished by the relative abundance of specific OTUs (Figure 1). The number assigned to each CST indicates the overall frequency of occurrence at each respective site, with CST 1 being the most frequent. Based on OTU abundance and the sequence of progression of CSTs over time observed in each subject (Figure 2), we concluded that the CSTs consistently exhibited three properties: 1) they had highly dissimilar composition between different body sites, 2) they were associated with post menstrual age (PMA), gestational age (GA) at birth, and/or week of life (WOL) and 3) they demonstrated non-random patterns of co-occurrence such that the observation of a specific CST at a given body site was highly predictive of CSTs at other body sites.

**Figure 1.**
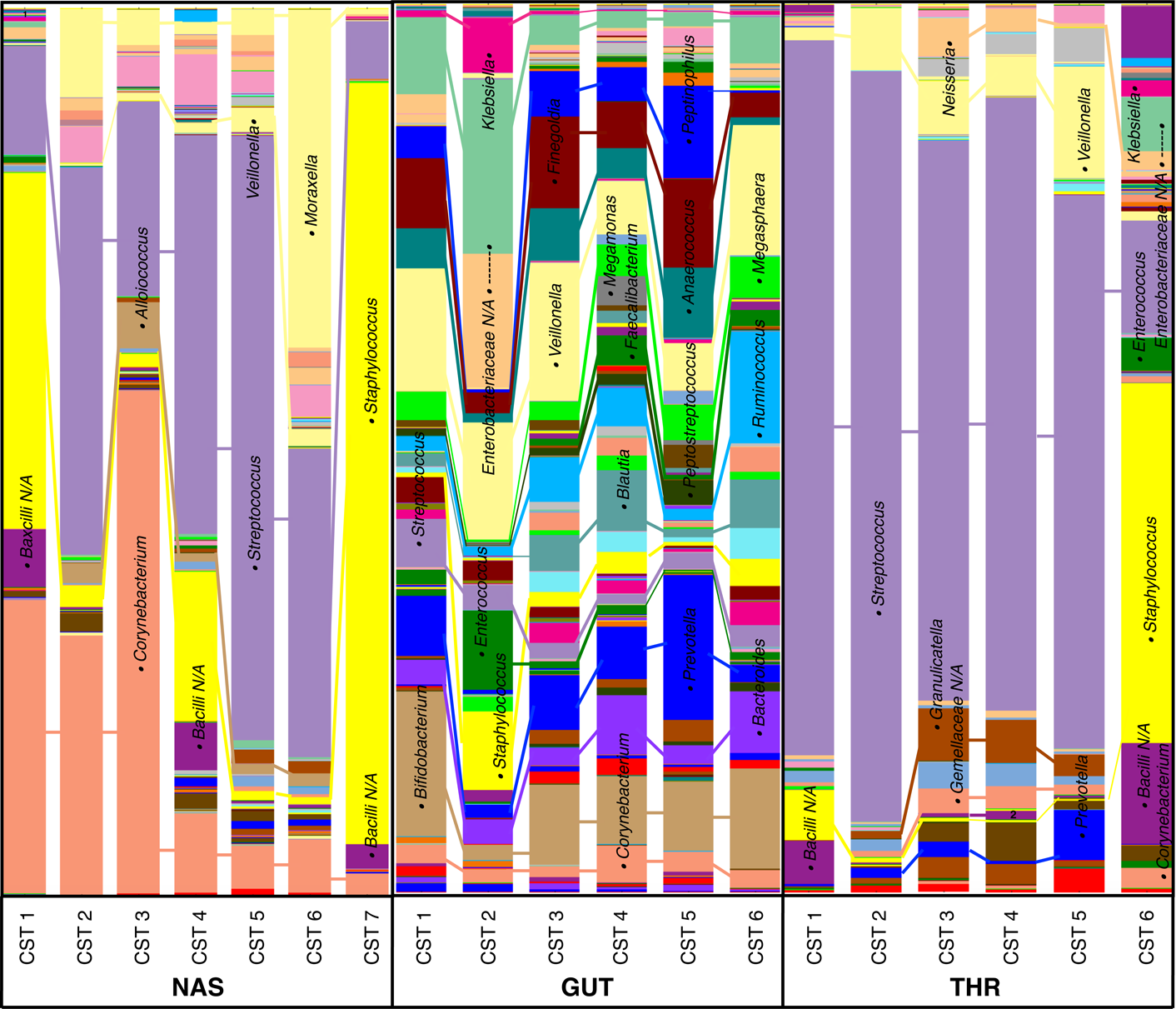
Composition of community state types (CSTs) of the nose (A), throat (B) and gut (C). Average composition of each CST was identified by Dirichlet Multinomial mixture (DMM) model-based clustering. Samples are grouped by the Dirichlet component that they represent, with each component corresponding to a CST, and the average composition of all samples in each CST group is represented. The height of each bar is equal, indicating that all total abundances are normalized to a constant sum. Within each bar, different colored bands correspond to different taxa, and the height of a given band is proportional to the average relative abundance of the corresponding taxon in the given CST. The top ten most abundant taxa within each body site are identified, with the closed circle flanking each taxa name positioned in the corresponding taxa in each bar. The composition of all samples is listed in Supplemental **Table 1**.

**Figure 2.**
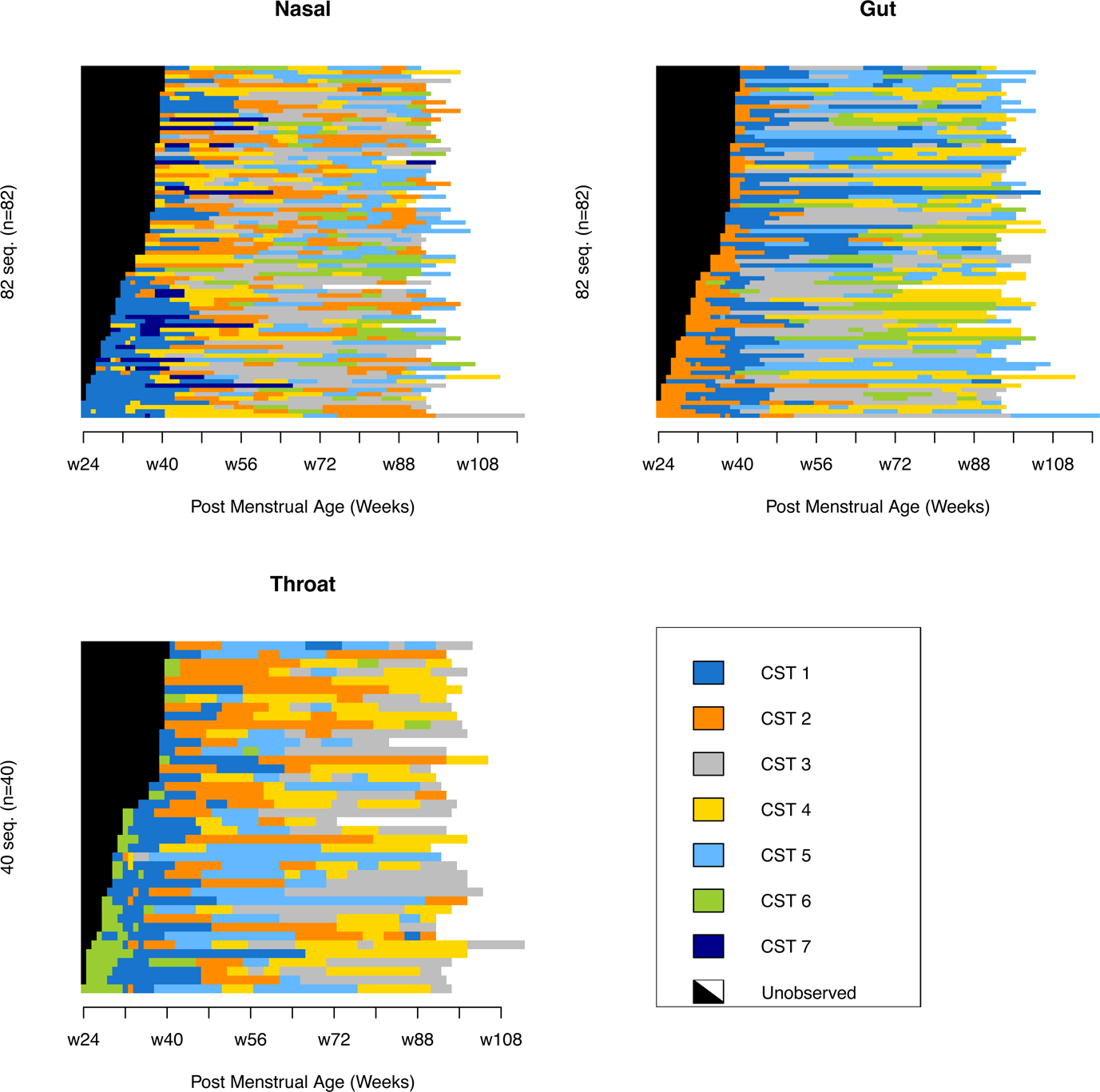
Sequence index plots indicate progression of community state types (CSTs) over time for each subject. Subjects are stratified along the y-axis and sorted in descending order by gestational age at birth. Post menstrual age (PMA) in weeks is indicated along the x-axis. The period of sampling for each individual is colored, with colors indicating the observed CST in a given time period. The time point of each observation is rounded down to the week in which the sample was taken and the surrounding period of time is colored according to the CST of the sample, with color changes occurring at the midpoint between consecutive samples in which different CSTs were observed. For each subject, the black region on the left indicates the period prior to birth and the white region on the right indicates the period after the last sample was taken. In all three body sites, strong temporal structure and ordered patterns of CST progression are evident. For example, CSTs 1, 2, and 6 are overrepresented during the period prior to 40 weeks PMA in the nose, gut, and throat, respectively.

### Microbiota composition of community state types

The multiple CSTs at different sites displayed a range of diversity and abundance of specific OTUs **(Figure 1)**. Notably, in all three body sites, the CST most frequently observed at the earliest time points had exceptionally high levels of *Staphylococcus*, which was much higher in samples of these earliest CST than in typical samples from any given body site. The throat CSTs were the least diverse, with types 1 through 5 dominated by *Streptococcus* (62%–85%) and type 6 by *Staphylococcus* (41%). With the exception of type 6, throat CSTs were differentiated from one another by lowly abundant but ubiquitous taxa, including *Veillonella*, *Granulicatella*, *Gemellaceae*, *Prevotella*, and a *Bacilli* that was not identifiable to the genus level by our methods. Nasal CSTs were substantially more diverse, with *Streptococcus* and *Corynebacterium* ranging from 5% to greater than 50% abundance across all CSTs. *Staphylococcus* (40%) and *Corynebacterium* (33%) comprised the majority of the community in nasal CST 1. Nasal CSTs 2 through 6 contained significant levels of *Streptococcus* (22%-68%), with types 2-5 differentiated primarily by lowly abundant taxa and the abundance of *Corynebacterium*, and type 6 distinguished by highly elevated levels of *Moraxella* (38%). Nasal CST 7 was dominated by *Staphylococcus* (86%). The community types of the gut were the most diverse of all three body sites and were consistently populated with *Enterobacteriaceae*, *Veillonella*, *Ruminococcus*, *Streptococcus*, *Prevotella*, *Bacteroides*, and *Bifidobacterium* at mean relative abundances greater than one percent. Individual gut CSTs were differentiated by the abundance of *Bifidobacterium* in CST 1 (16%); high abundance of *Staphylococcus* (9%), *Klebsiella* (6%), Enterobacteriaceae (35%), and *Enterococcus* (9%) and the absence of *Finegoldia* in CST 2 and of *Peptoniphilus* in CSTs 2 and 6. Type 6 was notable for elevated levels of *Ruminococcus* (13%). Gut CSTs 3, 4, and 5 were distinguished from one another primarily by lowly abundant taxa, but more prominent differentiating features included elevated abundance of *Corynebacterium* (4%), *Megamonas* (3%), and *Faecalibacterium* (3%) in CST 4; elevated levels of *Veillonella* (16%) and *Bifidobacterium* (9%) in CST 3; and elevated levels of *Prevotella* (16%), *Peptoniphilus* (10%), and *Peptostreptococcus* (3%) in CST 5. The average abundance of all genera in each CST from each body site is indicated in Figure 1 and listed in **Supplemental Table 1**.

### Community state type occurrence across chronological and developmental time

We next examined the association of CST occurrence with PMA in the 82 pre- and full-term infants. Patterns of temporal progression appear to be generally shared across individuals, with a majority of infants manifesting most community state types in their first year, in a similar sequence and at similar ages. These properties allow us to use CSTs to describe the ecological succession and summarize the development of the infant gut and respiratory microbiota in a conceptually and analytically tractable way. As described below, the occurrence of all throat, nasal and gut CSTs were strongly associated with PMA in both pre- and full-term infants **(Supplemental Figure 1)**.

*Throat CST and PMA.* Throat CST 6, being comprised in large part of *Staphylococcus*, is the only throat CST not dominated by *Streptococcus* and is typically the first CST to be observed, especially in pre-term infants. CST 6 quickly gives way to CST 1, which makes up the majority of observations in the first 2-3 months of life and is achieved more quickly but sustained for less time in full-term subjects than pre-terms. CST 2 is the next to emerge, reaching peak abundance in the first six months of life, manifesting earlier and with greater frequency in full-term than pre-term subjects and being the most common full-term CST from 42-65 weeks PMA. Pre-term subjects exhibit CST 5 more frequently and persistently than full-term subjects, which peaks in prevalence at ~25 weeks of life. CST 3 appears shortly after birth, several months before CST 4; both at low frequency initially and increasing in prevalence for the duration of the sampling period, accounting for a majority of samples beyond 70 weeks PMA and distinguishable only by lowly abundant and rare taxa.

*Nasal CST and PMA.* Pre-term infants overwhelmingly begin life in nasal CST 1, which is characterized by a high abundance of *Staphylococcus* and *Corynebacterium* and observed in fewer than 50% of full-term infants. The prevalence of CST 1 is reduced after 40 weeks and it is completely absent by 55 weeks PMA. CST 4, which is dominated by *Streptococcus*, is observed in the majority of subjects and appears at nearly all time points, but is most prevalent at 37-60 weeks PMA in both pre- and full-term infants. CSTs 2 and 3, which are dominated by *Corynebacterium* and *Streptococcus,* are common at all time points after 40 weeks PMA and comprise the majority of all samples beyond 60 weeks PMA. CST 5, which is characterized by a high abundance of *Streptococcus*, first appears at 50-60 weeks PMA and is observed more frequently in full-term subjects. CSTs 6 and 7, which are rare and are distinguished by high levels of *Moraxella* and *Staphylococcus*, respectively, are sporadically observed after 42 weeks PMA with CST 7 being more common prior to 60 weeks PMA and CST 6 being more common for the remainder of the period of observation.

*Gut CST and PMA*. Gut CST 2 comprised the majority of pre-term samples prior to 40 weeks PMA and full-term samples prior to 42 weeks PMA, but quickly gives way to CST 1 in most subjects. CSTs 1 and 3 account for the majority of observations through 60 weeks PMA, with CST 3 being more common in pre-term infants and typically occurring after CST 1. CSTs 4 and 5 were prevalent beyond 42 weeks PMA and accounted for the majority of samples by one year in both pre- and full-term infants, with CST 4 being especially common later on. Community state type 6 is observed in a minority of pre- and full-term subjects after 42 weeks PMA and peaks in abundance around one year of post-natal age, with no significant association with GA at birth.

Overall, a strong temporal structure and ordered progression of CSTs relative to PMA at all three body sites is apparent. While no single CST is observed in all 82 infants, common patterns of sequential CST occurrence at each body site reveal canonical orderings, with the throat progressing through CSTs 6, 1, 2, 5, 4 and 3; nasal through CSTs 1, 7, 4, 2, 3, 6 and 5; and gut through CSTs 2, 1, 3, 6, 4 and 5. However, pre- and full-term infants are initially colonized by distinct CSTs, with CSTs 1, 2 and 6 overrepresented prior to 40 weeks PMA in the nose, gut and throat, respectively. Furthermore, individual infants matched by PMA transition through CSTs at different rates, which suggest factors other than age regulate microbiota progression.

### Correlations between community state type and PMA in pre- and full-term infants

In order to elucidate the relationship between time and the progression of community types through each body site we further examined the associations between CSTs and time, which can be measured developmentally as PMA or postnatally with WOL. We first fit smoothed curves of the probability of being in a given CST against WOL and GA at birth (Figure 3) and the probability of being in a given CST against PMA and GA at birth **(Supplemental Figure 2)** which revealed several canonical temporal trends in the occurrence of CSTs. The chronological CSTs, a minority of all CSTs identified, showed no substantive difference over the first year of life between pre- and full-term infants (e.g. throat CSTs 3 and 4), indicating that the week of life, a proxy for environmental exposure, drives the occurrence of these community types. In contrast, the idiosyncratic CSTs appear to be endemic markers of GA at birth, with persistent differences in the probability of occurrence between pre-term and full-term infants for the first year of life (e.g. nasal CSTs 1 and 5, throat CST 6). Lastly, the convergent CSTs showed increased probabilities of occurrence at earlier post-natal ages in full-term infants, but CST occurrence probability in pre-term infants reached parity after a post-natal interval proportional to their prematurity (inversely proportional to their GA at birth).

**Figure 3.**
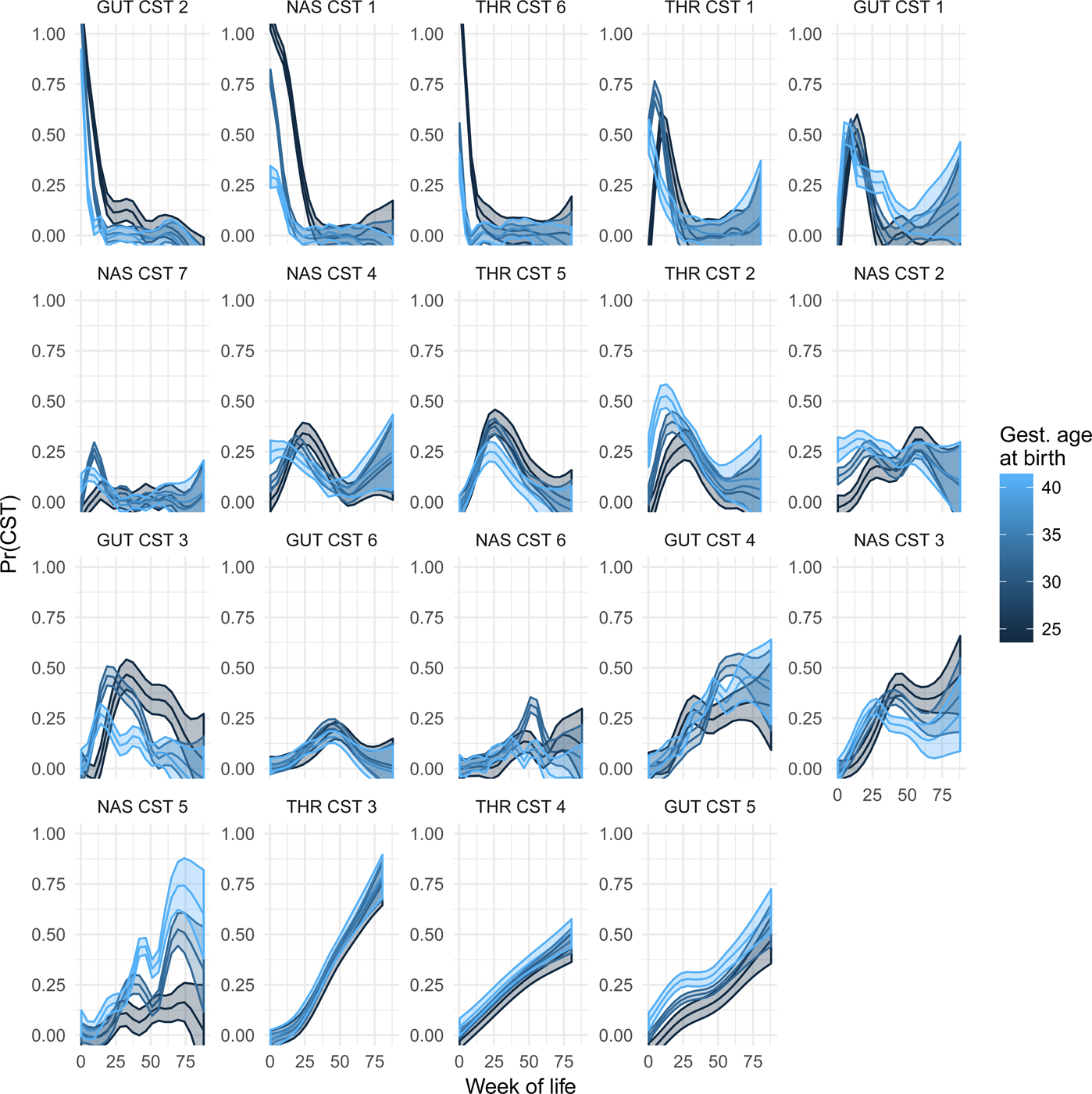
Associations between community state type membership and time. The posterior probability of membership to each CST (y-axis) is plotted over weeks of life (x-axis), estimated as a non-parametric function of week of life and gestational age at birth. The CSTs are sorted by post-menstrual age at which they achieve maximal probability of occurrence.

We then constructed a single index model of age for each CST, which fit the probability of observing the CST as a function of some combination of GA at birth and WOL **(Supplemental Figure 3A)**. These single index models confirmed the temporal trends characteristic of the three CST types described above and allowed us to quantify the differential effects of time spent pre- and postnatally, with respect to the probability of manifesting a given CST. These models were consistent with the trends described above, with three basic patterns being observed: CSTs for which GA at birth was not significant, but week of life was (chronological); CSTs for which a single index could not be well fit, indicating that no amount of time since birth could make up for disparities in GA at birth (idiosyncratic); and CSTs for which GA at birth and week of life were both significant, such that pre-term infants could catch up to full-term infants after some period of time (convergent). In this latter category of CSTs, the pre-term infants typically catch up at a rate of one week of life per week pre-term (i.e. gut CST 2, throat CST 1, 2 and 5), implicating developmental age (PMA) as the primary temporal correlate **(Supplemental Figure 3B)**.

### Associations between community types and composition across body sites

The tendency for CSTs in each body site to depend on PMA and postnatal age suggested potential relationships between CSTs across body sites. As expected, the co-occurrence patterns of CSTs across body sites were significantly non-random, as assessed by a Chi-squared test (p-value << 0.001). To further explore these associations, we calculated the pairwise correlations between CSTs observed at each site (Figure 4). Again, co-occurrence patterns between sites was highly significant, suggesting that the observation of a given CST at one body site is highly predictive of the CST at other body sites. The greatest degree of CST correlation between body sites was observed among nasal CST 1, gut CST 2, and throat CSTs 1 and 6, for which all cross-body site pairs were positively correlated.

**Figure 4.**
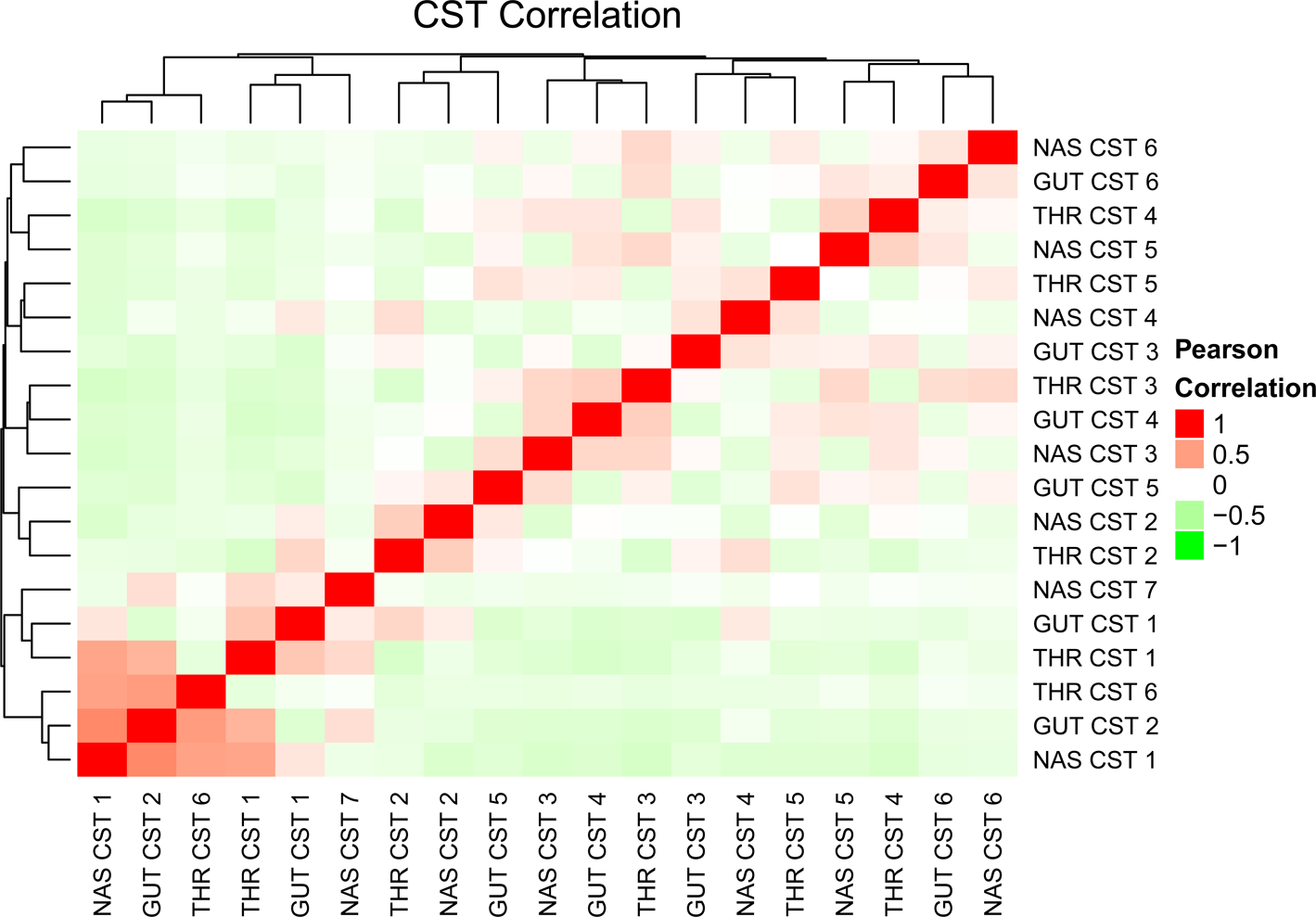
Pairwise correlations between community state types (CSTs) at different body sites. CSTs on the x- and y-axes are identified by body site (NAS: nasal, THR: throat, GUT: gut) and type. Each cell represents the Pearson sample correlation of CST membership probability across body sites in the same individual. Red hued cells correspond to positive correlation coefficients. CST co-occurrence is non-random with the CST of one body site highly predictive of the CSTs of the other two body sites.

Given the strong associations at all body sites between CST occurrence and infant developmental and chronological age, correlations across body sites are expected. In order to control for these factors and identify potential associations arising from direct or indirect interactions across sites, we further assessed the associations between the CST of each body site and the microbiota composition of the other body sites using a series of linear regression models. As predictors for the abundance of each taxon in a given body site, we first used mode of delivery, GA at birth, and day of life (which was modeled with a natural spline to allow for nonlinear effects), as well as a per-subject random effect to account for repeated sampling of the same individuals. We then added additional predictive terms for the CSTs of the other body sites and refit the models. Because we sought to identify the relationships across body sites that could not be explained by infant maturity alone, we called significant only those associations between taxon abundance and remote CST for which inclusion of the remote CSTs as terms in the model significantly improved its explanatory power (see methods). We identified significant associations across all pairs of body sites **(Supplemental Tables 1 and 2)**, with the most significant associations identified between CSTs of the nose and gut and taxa in the throat. Fewer associations were significant between the gut and nose. Within each body site, certain taxa were uniquely associated with the CST of only one of the other body sites, while other taxa exhibited significant associations with the CST of both of the other two body sites.

A number of taxa had significant associations with CSTs of other body sites at which the taxa themselves were not observed, ruling out direct exchange of these bacteria as the sole explanation for the associations. These taxa included *Bacteroides ovatus*, *Clostridium perfringens*, *Actinobaculum, Faecalibacterium sp.* (**Supplemental Table 2).** Instead these associations were consistent with the presence of bacteria in one site impacting, or being impacted by, development of microbiota at another site through indirect physiological or metabolic mechanisms. In order to assess cases where taxa were present in both associated sites, including *Viellonella*, *Prevotella* and *Dorea*, we tested the OTU residuals for correlation after adjusting for PMA with a spline and each subject with a mixed effect. There were approximately fifty shared OTUs between each pair of sites, which on average were positively correlated for each site pair. The strongest correlation between shared OTUs was between the nose and throat, followed by the throat and gut, followed by the nose and gut **(Supplemental Figure 4).**

The associations between each set of CSTs and specific taxa **(Supplemental Tables 1 and 2)** was visualized in two ways. First, as a bipartite graph (Figure 5A-C) in which each site’s most taxonomically specific significant taxa were connected to the CSTs of the distal body sites to which they had significant associations at a FDR of 10%. Edge color indicates the direction and significance of the association, either as a decrease or increase in abundance when the associated remote CST is observed. Second, as a volcano plot (Figure 5D), which indicates the significance (F-test p-value) and the magnitude of the increase in explanatory power (R^2^) when the CSTs of distal body sites are added to the regression models that include as covariates mode of delivery, GA at birth, day of life (as a natural spline), and a per subject random effect, and taxon abundance as the outcome. We identified 140 unique taxa with significant cross-body site associations; 59 in the gut, 61 in the nose, and 66 in the throat, with some taxa being significant in multiple body sites where they occurred. In the gut, 15 taxa were significantly associated with CSTs in both the throat and nose, 11 taxa in the nose were associated with both gut and throat CSTs, and 23 taxa in the throat were associated with both nose and gut CSTs. Among taxa present in the gut, the largest numbers of associations were identified with nasal CST 1 and throat CST 2, with 38 and 13 taxa respectively, and the single most significant association was between *Bacteroides ovatus* and throat CST 2 (Figures 5B and D). Notably, *B. ovatus* was not identified in throat samples but was present in 1% of nasal samples at a low (<1%) abundance. Among taxa present in the nose, the largest numbers of associations were identified with gut CST 1 and throat CST 2, with 26 and 29 taxa respectively, and the single most significant association was between an OTU of *Prevotella* and throat CST 2 (Figures 5A and D). Among taxa present in the throat, the largest numbers of associations were identified with nasal CST 1 and gut CST 1, with 22 and 36 taxa respectively, and the single most significant association was between *Prevotella pallens* and nasal CST 6 (Figures 5C and D). In the gut, an OTU of *Dorea* exhibited the most significant associations, with six CSTs from the nose and throat found to be significant. In the nose, an OTU of *Veillonella* had the most associations, with seven CSTs from the gut and the throat. In the throat, the *Lachnospiraceae* family and an OTU of *Veillonella* had the most associations, each with six CSTs from the nose and gut.

**Figure 5.**
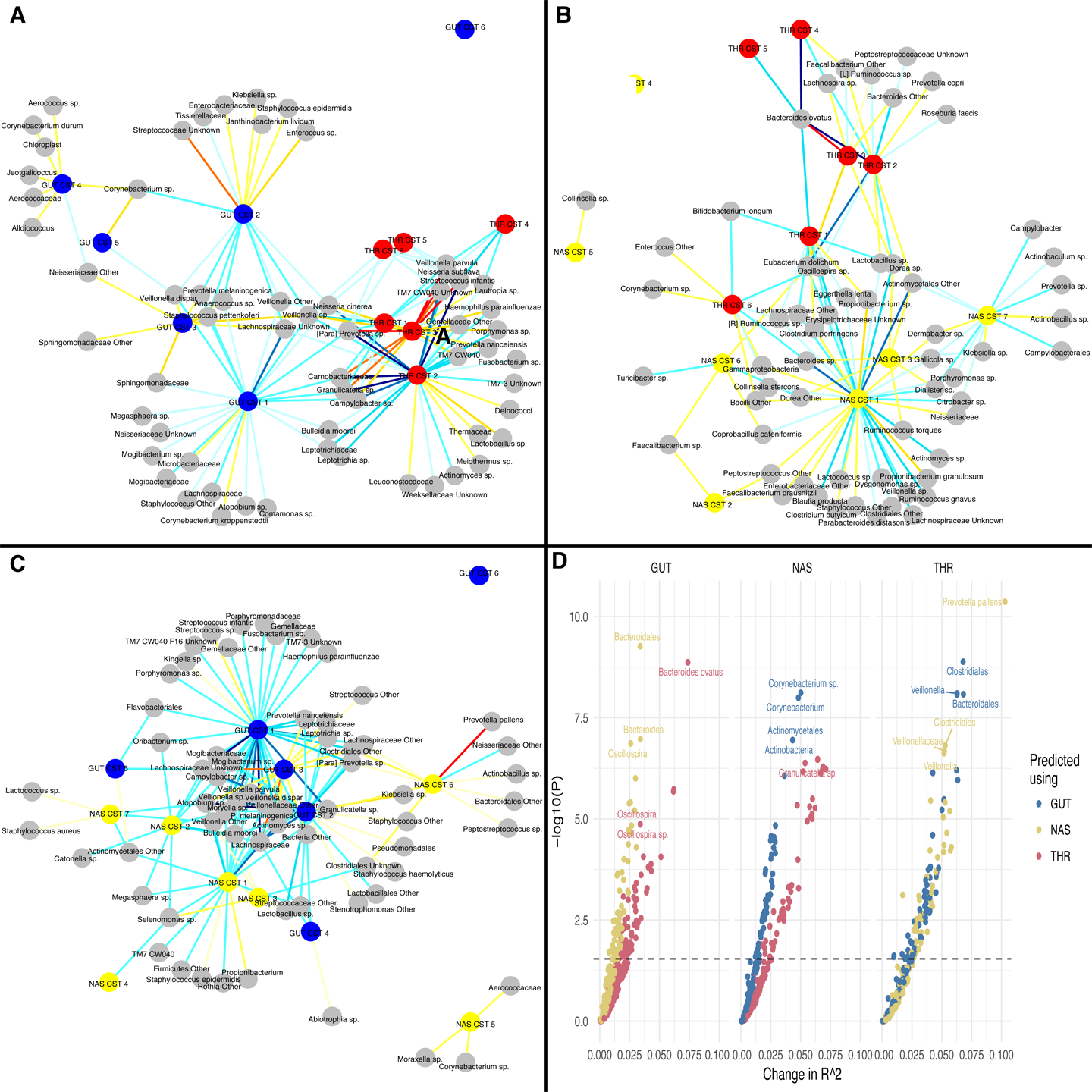
Significant associations between taxa abundance and community state type (CST) across body sites. A bipartite graph was used to visualize the associations between CSTs and taxa at a distal body site [**(A)**: nasal, **(B)**: gut, **(C)**: throat], with significant associations at a false discovery rate (FDR) of 10%. Edges indicate significant associations with color marking the direction of the effect (red: increase in abundance, blue: decrease in abundance). Color shade corresponds to level of significance, with lighter colors being less significant. Nodes are positioned using a force-directed layout, which places taxa or CSTs with similar patterns of significant associations near each other while attempting to optimize readability and limit overlap. **(D)** Relationships between taxa abundance and CSTs was also visualized using a volcano plot, with improvement in explanatory power (R^2^) conferred by the inclusion of CSTs in the model on the x-axis and -log10 p-values of the model improvement on the y-axis. With individual taxa in each body site as the outcome (subplots NAS, GUT and THR), a linear regression model was fit using the with and without CSTs of the other body sites as covariates, controlling for gestational age at birth, day of life, mode of delivery, and subject-level random effects. Full models (including all CSTs) were tested against null models (excluding the CSTs of the other body sites in turn) with a series of F-tests.

### The microbiome is canonically correlated across body sites following time and space

These observations prompted us to assess the extent to which the OTU composition of each body site over time can be explained as a function of the OTU composition of the other body sites, without the dimension reduction associated with using DMMs and CST classification. We again paired microbiome samples from different body sites that were acquired at the same visit for each participant, resulting in nasal-gut, nasal-throat and rectal-throat site pairs. We then assessed the correlation of taxa between body sites directly using canonical correlation analysis (CCA), which transforms two sets of multivariate observations into a series of *canonical correlates* (weighted averages) that maximize the score cross-correlations, quantifying the extent to which two sets of multivariate observations are correlated. Using a cross-validation scheme that held out blocks of individuals, we found in validation data that the subspace cross-correlations varied from 0.75 (nasal-throat) to 0.6 (gut-throat), for the first canonical coordinate **(CC) (Supplemental Figure 5)**. Since we previously established that many OTUs vary as a function of time, we anticipated that temporal variation was responsible for much of this correlation. As expected, after adjusting for time by regressing out PMA in each site with a 14 degree-of-freedom spline, the time-stabilized correlations were attenuated in all site pairs, but still significantly different from zero in the nasal-gut and nasal-throat pairs **(SupplementalFigure 5)**.

## DISCUSSION

Development of the infant microbiome landscape is a critical factor in overall infant development and long-term health. In this study, we examined the progression of and spatiotemporal interactions between the gut and respiratory microbiota in pre- and full-term infants. Knowledge of inter-site interactions between these anatomical niches forms the basis for understanding microbiota perturbations in sick infants and potential alterations in the microbiota of other sites. We combined a community state type framework with longitudinal modeling to identify significant associations between microbiota across the nose, throat, and gut during early life development in pre- and full-term infants. We demonstrate that the abundance of specific taxa in one body site exhibited strong associations with the community types of the other body sites. Using a natural spline function to control for time and linear regression to control for GA at birth, mode of delivery, and within subject correlation, we found that incorporating the community types of the other body sites significantly improved the explanatory power of our model for the abundances of 140 unique taxa, many of which were present in only one member of a pair of associated body sites, ruling out the possibility of direct transmission as the mechanism of association. We then performed canonical correlation analysis to validate the explanatory power of the composition of each body site for every other body site. While the effect of time accounted for the majority of the canonical correlation between sites, a significant degree of correlation was observed even after controlling for temporal effects. These observations suggest a potential systemic coordination of microbial abundance and distribution across the infant microbiota landscape during early life development.

Early infant microbiota studies have focused on single anatomical sites, such as the gut and respiratory microbiota, with unique communities and distinct functions. Studies on pre- and full-term infant gut microbiota have shown that postnatal microbial colonization initiates maturation of infant intestinal structures and mediates development of the immune system through interactions with gut epithelial, immune effector and mucus producing cells [24-26]. Deficiency in colonization of pre-term infant gut microbiota has been associated with delays in immune development, alterations in host metabolism and inflammatory diseases such as necrotizing enterocolitis (NEC) [11, 27-31]. Longitudinal studies with pre-term infants have shown that the gut microbiota develops in a series of phases associated with postmenstrual age (PMA), more so than post-natal age, suggesting possible coordination between microbiota maturation and functional differentiation of the gut epithelium at defined stages of infant development [32]. Recent reports on neonatal respiratory microbiota have identified similar interactions of microbiota with mucosal epithelial and immune cells and an association with respiratory tract infections and chronic lung disease of prematurity [33-36]. However, most respiratory microbiome studies have focused on a limited number of samples from full-term infants. The influence of the respiratory microbiota on lung immunity and respiratory diseases in high risk pre-term infants underscores the need to better understand initial microbial colonization and temporal dynamics of respiratory microbiota through longitudinal studies as we describe here.

The gut and respiratory tracts in infants share the same embryonic origin, with mucosal surfaces composed of columnar epithelial cells that sense the commensal microbiota and in turn shape local and systemic immunity as infants mature and as a function of PMA [24-26, 35, 37]. Changes in infant gut and respiratory microbiota that occur as a result of diet, antibiotics, therapeutics and environmental exposures in the NICU are likely to influence the microbiota at both sites [32, 38, 39]. The effect of these changes can be illustrated by antibiotic induced alterations of neonatal gut microbiota during the crucial early postnatal period of immune competence, which increase the risk of developing allergic airway disease and other atopies in subsequent childhood [40-42]. In adults, common chronic lung diseases, such as asthma and chronic obstructive pulmonary disease (COPD) often coincide with inflammatory bowel disease (IBD) and other chronic gastrointestinal syndromes [37, 43, 44]. The occurrence of these chronic lung diseases is accompanied by functional and structural changes in the intestinal mucosa and increased intestinal permeability, suggesting that interactions between these two distal sites through the gut-respiratory axis impact adult health and disease [43, 45]. These gut-respiratory interactions likely function on several levels, ranging from direct transfer of bacteria between these sites through reflux and aspiration to indirect effects from bacterial metabolic products or mucosal immune responses common to both the gut and respiratory tract [33, 35, 37, 46]. Taken together, these observations of common developmental origins for the gut and respiratory tracts as well as inflammatory diseases that affect both sites, support potential systemic mechanisms that coordinate microbiota development at these distal sites in infants.

The microbiota samples for our study were collected as gut, nasal and throat swabs from pre- and full-term infants. In a previous study, we established the taxonomic similarity of infant gut microbiota samples collected either as rectal swabs or from fecal material on a diaper [32]. When evaluating respiratory samples for this study and their relatedness to lung microbiota, we first considered potential acquisition routes for respiratory microbiota. The lung microbiota in healthy individuals is acquired by direct mucosal dispersion and micro-aspiration from the upper respiratory tract (URT) [47, 48]. The microbiota in these sites are taxonomically similar, albeit with differences within the URT subcompartments (nasal cavity, nasopharynx, oropharynx and trachea) and lungs a result of cellular and physiological features, such as oxygen and carbon dioxide tension, pH, humidity and temperature that distinguish these environments and select for particular taxa [47-49]. The nasopharynx and oropharynx are the primary sources of lung microbiota in infants, likely due to the anatomy of the infant URT and increased production of nasal secretions, both of which enhance dispersal of microbiota to the lungs [50, 51]. With the infant nasopharynx and oropharynx, a primary source of colonizing infant lung microbiota, the nasal and throat samples used in our study as representative proxies of the neonate lung microbiota identified significant associations of taxa and CSTs within the gut-respiratory axis. The orthogonal variation of the nasal and throat microbiota is noteworthy, suggesting that additional analysis of both sites will provide unique insights into gut-respiratory interactions.

In our initial observations of the CST microbiota content relative to PMA, we noted that*Staphylococcus* was the most abundant taxa in the first CST of all three body sites (Figure 1 and **Supplementary Figure 1)**. Subsequent CSTs in all three sites demonstrated a rapid decrease in *Staphylococcus* abundance, which was progressively replaced by site specific taxa with cellular and metabolic capabilities required for adaptation to the developing host site and interaction with the colonizing microbiota. Previous studies of infant gut microbiota identified *Staphylococcus* as an early microbiota colonizer, with abundance determined by nutrition and mode of delivery [8, 52]. With a metabolism biased towards carbohydrate metabolism, emerging data suggests the potential for a strong impact of *Staphylococcus* on disease programming and obesity in later life [53-55]. *In vitro* and *in vivo* animal experiments assessing transcriptomic and phenotypic responses of *S. aureus* to microbiota partners have revealed mechanisms that modulate metabolism, virulence and survival in a multi-species bacterial community [56-58]. Similar experimental approaches to study interactions between members of the microbiota are needed to assess the mechanistic foundation of microbiota associations identified through computational means.

Identification of taxa having significant associations with CSTs in other body sites, although not present in the site with which they were associated, suggested that underlying immune or metabolic mechanisms mediate microbiota development between the distal gut and respiratory sites. One hundred forty unique taxa were identified with significant cross-body site associations in the nose, throat and gut at a FDR of 10% (Figure 5, Supplementary Tables 1 and 2). A small number of taxa were significant in multiple body sites, with the most significant associations identified between *B. ovatus* in the gut and throat CST 2. A plausible basis for association with distal CSTs can be proposed for *B. ovatus* and other taxa. Evidence that *B.ovatus*, a gut symbiont, digests polysaccharides in the gut as a carbon source for other members of the *Bacteroides* genus, places it at the center of cooperative ecosystem that is likely a central factor for gut microbiota functions and potential interactions with microbiota at distal sites [59-62]. Production of small chain fatty acids (SCFA) produced by *Bacteroides* and other enteric bacteria have been shown to profoundly affect both mucosal and systemic antibody responses [63]. Furthermore, increased abundance of *B. ovatus* in the gut has been associated with systemic autoimmune diseases and IBD, a disorder linked to respiratory diseases as described above [64]. Overall, the identified taxa-CST associations have the potential to effect gut-respiratory crosstalk through production of bacterial metabolites and ligands. In turn, dysbiosis of the gut microbiota can be anticipated to affect dynamics of respiratory microbiota as well as systemic metabolic and immune responses [37].

In our examination of the associations between CSTs at each body site and time as measured developmentally by PMA or chronologically by WOL, we identified several canonical temporal trends in the occurrence of CSTs that were confirmed using single index models **(Supplemental Figure 3)**. Three groups of CST patterns were identified: 1) *chronological*, for which the only factor associated with manifestation of the community type was WOL, 2) *idiosyncratic*, for which there was a persistent disparity in occurrence patterns associated with GA at birth and 3) *convergent*, for which there is a temporal off-set between pre-term and full-term infants initially, but an eventual convergence. These results suggested that the assembly of infant microbiota has complex relationships with time and development, with the manifestation of certain community structures depending on developmental/gestational age, exposure/day of life or a combination of both, while others being permanently and persistently influenced by pre-term birth. Following observations made by other groups, we find that the strength of associations between microbial habitats is proportional to their proximity within the host [16]. The sites that are nearest to one another, such as the nose and throat, have the highest time-stabilized correlation, and are more likely to share species in the canonical coordinate (CCA) site loadings. Distal pairs, such as the nose and gut have lower canonical correlations, with greater heterogeneity in the CCA loadings. In other words, body sites that are closer together have more microbial taxa in common and exhibit stronger associations between their microbiota composition than sites that are farther apart. These findings demonstrate that significant canonical correlations exist between the composition of microbial communities across body sites which cannot be entirely attributed to each body site’s independent temporal progression or to the repeated sampling of the same individuals.

## Conclusion

Understanding the variation between and within subjects, conditions, and over time as the manifestation of distinct community types provides an attractive conceptual and analytical framework for studying the microbiome [65]. While the extent to which community types are discrete or simply represent dense locations on a continuous gradient appears to vary depending on the conditions being sampled and the definition of “community type”, both the theoretical basis for community types and the utility of a community type-based framework has been established [65-73]. Significantly, the underlying network structure observed in microbiota gives rise to stable community types and a community type-based model is appealing in that it is analytically tractable and circumvents many of the complications associated with high dimensional compositional data. Furthermore, the utility of community types can be extended to the identification of associations across body sites. Ding and Schloss used Dirichlet multinomial mixture modeling to define canonical community types within 18 adult body sites independently and then demonstrated that while the community types across different body sites were dissimilar in composition, they were predictive of one another [66]. The occurrence of community types simultaneously in different body sites was highly non-random, suggesting an unknown mechanism of coordination or interaction acting at a distance. However, the community type framework may mask important biological variability and lack power to detect specific taxa that serve as superior phenotypic biomarkers [65]. The approaches taken in our work reported here largely mitigate these shortcomings, by employing a sampling scheme that was dense and evenly distributed over gestational ages at birth, week of life, and modes of delivery, thereby making it unlikely that apparent community clusters are the result of a failure to observe intermediate points along a gradient. Our subsampling procedure to determine the number of clusters yielded a robust and parsimonious description of the data. Community types were not confounded within individuals, but shifted in type over the period of observation for each individual and were seen across a plurality of individuals. In this setting, where we sought to characterize the development of respiratory and gastrointestinal microbiota over the first year of life, community types have provided a high-level description of the state and progression which facilitated the interrogation of associations of CSTs with developmental age (PMA) and post-natal chronological age (WOL).

In summary, new clinical strategies for establishing and maintaining a homeostatic microbiota are needed for neonates at risk for gut and respiratory dysfunction and immune deficiencies. A greater understanding of infant respiratory microbiota colonization, interactions between the respiratory and gut microbiota and possible developmental coordination between the two body sites are crucial steps in that direction. Our results demonstrate the existence of a host-wide network of associations between microbiota. The fact that these associations cannot be entirely explained by time, subject, or direct exchange of bacteria suggest unobserved factors mediating microbial dynamics and associations between microbiota across environments and at substantial distances. To our knowledge, these observations directly implicate, for the first time, a body-wide systemic mechanism coordinating the abundance and distribution of microbiota during early life development. The methods employed here may facilitate future efforts to evaluate disease, developmental maturity, therapeutic interventions, and dynamic interactions between multiple microbial communities and host systems.

## METHODS

### Clinical methods

All study procedures were approved by the University of Rochester Medical Center (URMC) Internal Review Board (IRB) (Protocol # RPRC00045470) and all subject’s caregivers provided informed consent. We sampled 1,079 gut (279 from NICU and 800 post-discharge), 1,013 nasal and (262 from NICU and 751 post-discharge) and 538 throat (172 from NICU and 366 post-discharge) microbiota samples longitudinally from 38 pre-term and 44 full-term infants. The Infants included in the study were from the University of Rochester Respiratory Pathogens Research Center PRISM study and cared for in the URMC Golisano Children’s Hospital Gosnell Family Neonatal Intensive Care Unit (NICU) or in the normal newborn nurseries and birthing centers. Fecal (rectal), nasal, and throat material was collected from pre-term infants from 23 to 37 weeks GA at birth (GAB) weekly until hospital discharge and then monthly through one year of age, adjusted for prematurity. Rectal, nasal and throat samples were collected from full-term infants at enrollment and monthly through one year. Each fecal sample was collected by inserting a sterile, normal saline moistened, Copan^®^ flocked nylon swab (Copan Diagnostics, Murrieta, CA) beyond the sphincter into the rectum and twirling prior to removal. Nasal and throat samples were similarly collected by inserting and twirling a sterile, moistened swab into the throat or anterior nostril. Each swab was then immediately placed into sterile buffered saline and stored at 4° C for no more than 4 hours. Samples were processed daily, which involved extraction of the fecal, nasal and throat material from the swabs in a sterile environment and immediately frozen at −80° C until DNA extraction. All sampling swabs, plasticware, buffers and reagents used for sample collection and extraction of nucleic acids were sterile and UV-irradiated to insure no contamination from sources outside of the infant.

### Genomic DNA extraction

Total genomic DNA was extracted from the nose, throat and rectal samples using a modification of the ZymoBIOMICS^TM^ DNA Miniprep Kit (Zymo Research, Irvine, CA) and FastPrep mechanical lysis (MPBio, Solon, OH). 16S ribosomal DNA (rRNA) was amplified with Phusion High-Fidelity polymerase (Thermo Scientific, Waltham, MA) and dual indexed primers specific to the V3-V4 (319F: 5’ ACTCCTACGGGAGGCAGCAG 3’; 806R: 3’ ACTCCTACGGGAGGCAGCAG 5’) and V1-V3 (8F: 5’ AGAGTTTGATCCTGGCTCAG 3’; 534R: 3’ ATTACCGCGGCTGCTGG 5’) hypervariable regions [74]. Amplicons were pooled and paired-end sequenced on an Illumina MiSeq (Illumina, San Diego, CA) in the University of Rochester Genomics Research Center. Each sequencing run included: (1) positive controls consisting of a 1:5 mixture of *Staphylococcus aureus*, *Lactococcus lactis*, *Porphyromonas gingivalis*, *Streptococcus mutans*, and *Escherichia coli*; and (2) negative controls consisting of sterile saline.

### 16S rRNA sequence processing

Raw data from the Illumina MiSeq was first converted into FASTQ format 2×300 paired end sequence files using the bcl2fastq program, version 1.8.4, provided by Illumina. Format conversion was performed without de-multiplexing and the EAMMS algorithm was disabled. All other settings were default. Sequence processing and initial microbial composition analysis were performed with the Quantitative Insights into Microbial Ecology (QIIME) software package [75], version 1.9.1. Reads were multiplexed using a configuration described previously [74]. Briefly, for both reads in a pair, the first 12 bases were a barcode, which was followed by a primer, then a heterogeneity spacer, and then the target 16S rRNA sequence. Using a custom Python script, the barcodes from each read pair were removed, concatenated together, and stored in a separate file. Read pairs were assembled using fastq-join from the ea-utils package, requiring at least 40 bases of overlap for V3V4 sequences and 20 bases of overlap for V1V3 sequence, while allowing a maximum of 10% mismatched bases. Read pairs that could not be assembled were discarded. The concatenated barcode sequences were prepended to the corresponding assembled reads, and the resulting sequences were converted from FASTQ to FASTA and QUAL files for QIIME analysis. Barcodes, forward primer, spacer, and reverse primer sequences were removed during de-multiplexing. Reads containing more than four mismatches to the known primer sequences or more than three mismatches to all barcode sequences were excluded from subsequent processing and analysis. Assembled reads were truncated at the beginning of the first 30 base window with a mean Phred quality score of less than 20 or at the first ambiguous base, whichever came first. Resulting sequences shorter than 300 bases or containing a homopolymer longer than six bases were discarded. Operational taxonomic units (OTU) were picked using the reference-based USEARCH (version 5.2) [76] pipeline in QIIME, using the May 2013 release of the GreenGenes 99% OTU database as a closed reference [77, 78]. An indexed word length of 128 and otherwise default parameters were used with USEARCH. Chimera detection was performed *de novo* with UCHIME, using default parameters [76]. OTU clusters with less than four sequences were removed, and representative sequences used to make taxonomic assignments for each cluster were selected on the basis of abundance. The RDP Naïve Bayesian Classifier was used for taxonomic classification with the GreenGenes reference database, using a minimum confidence threshold of .85 and otherwise default parameters [79].

### CST inference with Dirichlet Multinomial Modeling (DMM)

The DMM model was fit using the R package DirichletMultinomial version 1.16.0, R version 3.3.3. Sample composition was represented using normalized counts for each of the most specific operational taxonomic units (OTUs) present in at least 5% of the samples from a given body site. Normalization was performed on a per sample basis by taking the relative abundance of each OTU (after removing OTUs present in less than 5% of samples) and multiplying by 5,000. Resulting non-integer counts were rounded down. In the DMM model, the number of Dirichlet components is a tuning parameter. For each body site, we used 10-fold random subsampling of 80% of the samples to assess uncertainty in model fit for one through ten components, with model fit being assessed as the Laplace approximation of the negative-log model evidence. We selected the number of components at each body site corresponding to the lower bound on the standard error of the model fit. We then fit complete models for each body site using all samples and the number of components selected, and used the resulting posterior probabilities to assign each sample to a community state type (CST) corresponding to a Dirichlet component. The CSTs observed in each subject and at each body site over time are represented in Figure 2, which was plotted using the TraMineR package, version 2.0-7. Color changes occur midway between consecutive samples of differing CSTs. Observation time points were quantized for plotting purposes only, and this was done by rounding down to the nearest whole week of post menstrual age.

### Generalized Additive modeling of CST

Generalized additive models (GAMs) were fit using R package mgcv for each CST and site. The probability of being in a CST was modeled (on the linear probability scale) as a smooth function of week-of-life and GA at birth, and a random effect for each individual. Formally, for each CST, for individual *i* at time *t*, we fit

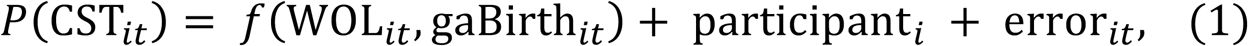

where *f*(WOL_*it*_, gaBirth_*it*_) is a smooth function of week of life and GA at birth, participant_*i*_ is a random intercept for each participant and error_*it*_ represents independent, homoscedastic noise. We plotted the fitted CST probability, under model (1) over a range of weeks-of-life for several representative gestational ages, and then compared this estimate to a single index model.

The single index model restricts the smooth function f in model (1) to seek a common “time” variable that accounts for both time spent inside, and outside the womb. Symbolically, we require *f* (WOL_*it*_,gaBirth_*it*_) = *f* (*a* · WOL_*it*_ + *b* · gaBirth_*it*_)). For instance, if a=1 and b=0 then GA has no effect on the CST trajectory, and when a=b then time spent inside the womb has the same effect on the probability of belonging to a CST as time spent outside the womb.

### CST and Taxa regression

We paired microbiome samples from different body sites that were acquired at the same visit for each participant. This generated 3 pairs of sites. The Nasal-Rectal sites had the greatest number of matched sample pairs, with 82 participants having 951 pairs of samples, while the Nasal-Throat sites had the fewest, with 40 participants having 483 sample pairs. The Rectal-Throat sites had 491 sample pairs. We applied arcsin sqrt-tranformation to stabilize the variance of relative abundance and then fit linear mixed effects models to the abundance using the CST of the other two sites as the primary variables of interest. We adjusted as potential confounders the mode of delivery, GA at birth and 14 degree-of-freedom spline for WOL. The subject ID served as a random intercept. Associations with the primary variables of interest were tested for all taxonomic levels and reported on the most specific taxon (or equally specific taxa) within a phylogenetic lineage. We report a Wald test for equality between the abundance in each CST and its grand mean abundance. Associations significant at 10% FDR, calculated per site, are shown as edges in Figure 5, which itself was generated using R packages GGally version 1.3.2 and network version 1.13.0. An overall test for association between a site and a taxon was derived by conducting an F-test of the model that dropped the CST of that site as predictors and the full model. The change in pseudo R^2^ reports the change in variance explained by the fixed effects in the null and full models [80].

### Canonical correlation analysis

The CCA function implemented in R base was used for canonical correlation analyses. We employed 10-fold cross validation in which we fit the CCA on each pair of sites on 9/10ths of the subjects, then calculated the subspace correlations on the 10% of withheld subjects. Two times the standard deviation of the held-out subspace correlation is shown in the shaded region of Supplemental Figure 5.

## LIST OF ABBREVIATIONS

GA: gestational age
PMA: premenstrual age
WOL: week of life
CST: community state type
OTU: operational taxonomic units
FDR: false discovery rate
NICU: neonatal intensive care unit

## DECLARATIONS

### Ethics approval and consent to participate

Written informed consent was obtained from parent or guardian of all participating infants. The institutional review board at the University of Rochester School of Medicine and Strong Memorial Hospital approved the study.

## CONSENT FOR PUBLICATION

Not applicable.

## AVAILABILITY OF DATA AND MATERIALS

All phenotypic data, 16S rRNA sequence reads and generated datasets is publically available through dbGaP accession phs001347.v1.p1.

## AUTHOR CONTRIBUTIONS

S.R.G., G.S.P., M.T.C., K.M.S., A.G. and A.M. designed the study. A.L.G., H.A.K. and H.H. collected and processed specimens. A.L.G. and H.A.K. sequenced and generated data. A.G., A.M., X.Q., J.J., H.Y., S.B. and J. H-W. analyzed data. A.G., A.M. and S.R.G. drafted the manuscript with revisions from G.S.P., M.T.C., K.M.S., S.B., A.R.F and D.J.T. All authors reviewed the final manuscript.

## ACKNOWLEDGEMENTS

We thank Deanna Maffett, Tanya Scalise, Gerry Lofthus, Melissa Bowman, Amy Murphy, Claire Wyman, Emily Fitzgerald and Elizabeth Werner for collection and recording of samples and clinical data. Microbiome sequencing in this study was completed by the University of Rochester Genomics Research Center (GRC).

## REFERENCES

1. Cho, I. and Blaser, M.J. The human microbiome: at the interface of health and disease. Nat Rev Genet, 2012. 13 (4): p. 260–70.

2. Backhed, F., Host responses to the human microbiome. Nutr Rev, 2012. 70 Suppl 1: p. S14–7.

3. La Rosa, P.S.,et al., Patterned progression of bacterial populations in the premature infant gut. Proc Natl Acad Sci U S A, 2014. 111 (34): p. 12522–7.

4. White, R.A., et al., Novel developmental analyses identify longitudinal patterns of early gut microbiota that affect infant growth. PLoS Comput Biol, 2013. 9 (5): p. e1003042.

5. Yatsunenko, T., et al., Human gut microbiome viewed across age and geography. Nature, 2012. 486(7402): p. 222–7.

6. Backhed, F., et al., Dynamics and Stabilization of the Human Gut Microbiome during the First Year of Life. Cell Host Microbe, 2015. 17(6): p. 852.

7. Costello, E.K., et al., Bacterial community variation in human body habitats across space and time. Science, 2009. 326(5960): p. 1694–7.

8. Dominguez-Bello, M.G., et al., Delivery mode shapes the acquisition and structure of the initial microbiota across multiple body habitats in newborns. Proc Natl Acad Sci U S A, 2010. 107(26): p. 11971–5.

9. Human Microbiome Project, C., Structure, function and diversity of the healthy human microbiome. Nature, 2012. 486(7402): p. 207–14.

10. Koenig, J.E., Succession of microbial consortia in the developing infant gut microbiome. Proceedings of the National Academy of Sciences, 2011(108): p. 4578–4585.

11. Costello, E.K., et al., Microbiome assembly across multiple body sites in low-birthweight infants. MBio, 2013. 4(6): p. e00782–13.

12. Chu, D.M., et al., Maturation of the infant microbiome community structure and function across multiple body sites and in relation to mode of delivery. Nat Med, 2017.

13. Gritz, E.C. and Bhandari, V. The human neonatal gut microbiome: a brief review. Front Pediatr, 2015. 3: p. 17.

14. Lim, E.S., et al., Early life dynamics of the human gut virome and bacterial microbiome in infants. Nat Med, 2015. 21(10): p. 1228–34.

15. Faust, K., et al., Cross-biome comparison of microbial association networks. Front Microbiol, 2015. 6: p. 1200.

16. Faust, K., et al., Microbial co-occurrence relationships in the human microbiome. PLoS Comput Biol, 2012. 8(7): p. e1002606.

17. Weiss, S., et al., Correlation detection strategies in microbial data sets vary widely in sensitivity and precision. ISME J, 2016. 10(7): p. 1669–81.

18. Kurtz, Z.D., et al., Sparse and compositionally robust inference of microbial ecological networks. PLoS Comput Biol, 2015. 11(5): p. e1004226.

19. Arrieta, M.C., et al., The intestinal microbiome in early life: health and disease. Front Immunol, 2014. 5: p. 427.

20. Borre, Y.E., et al., Microbiota and neurodevelopmental windows: implications for brain disorders. Trends Mol Med, 2014. 20(9): p. 509–18.

21. Renz, H., Brandtzaeg, P., and Hornef, M. The impact of perinatal immune development on mucosal homeostasis and chronic inflammation. Nat Rev Immunol, 2012. 12(1): p. 9–23.

22. Shukla, S.D., et al., Microbiome effects on immunity, health and disease in the lung. Clin Transl Immunology, 2017. 6(3): p. e133.

23. Holmes, I., Harris, K., and Quince, C. Dirichlet multinomial mixtures: generative models for microbial metagenomics. PLoS One, 2012. 7(2): p. e30126.

24. Hooper, L.V. and Gordon, J.I. Commensal host-bacterial relationships in the gut. Science, 2001. 292(5519): p. 1115–8.

25. Levy, O., Innate immunity of the newborn: basic mechanisms and clinical correlates. Nat Rev Immunol, 2007. 7(5): p. 379–90.

26. Sekirov, I. and Finlay, B.B. The role of the intestinal microbiota in enteric infection. J Physiol, 2009. 587(Pt 17): p. 4159–67.

27. Barron, L.K., et al., Independence of gut bacterial content and neonatal necrotizing enterocolitis severity. J Pediatr Surg, 2017. 52(6): p. 993–998.

28. Mai, V., et al., Fecal microbiota in premature infants prior to necrotizing enterocolitis. PLoS One, 2011. 6(6): p. e20647.

29. Morrow, A.L., et al., Early microbial and metabolomic signatures predict later onset of necrotizing enterocolitis in preterm infants. Microbiome, 2013. 1(1): p. 13.

30. Unger, S., et al., Gut microbiota of the very-low-birth-weight infant. Pediatr Res, 2015. 77(1–2): p. 205–13.

31. Warner, B.B. and Tarr, P.I. Necrotizing enterocolitis and preterm infant gut bacteria. Semin Fetal Neonatal Med, 2016. 21(6): p. 394–399.

32. Grier, A., et al., Impact of prematurity and nutrition on the developing gut microbiome and preterm infant growth. Microbiome, 2017. 5(1): p. 158.

33. Gallacher, D.J. and Kotecha, S. Respiratory Microbiome of New-Born Infants. Front Pediatr, 2016. 4: p. 10.

34. Gollwitzer, E.S. and Marsland, B.J. Microbiota abnormalities in inflammatory airway diseases - Potential for therapy. Pharmacol Ther, 2014. 141(1): p. 32–9.

35. Warner, B.B. and Hamvas, A. Lungs, microbes and the developing neonate. Neonatology, 2015. 107(4): p. 337–43.

36. Yun, Y., et al., Environmentally determined differences in the murine lung microbiota and their relation to alveolar architecture. PLoS One, 2014. 9(12): p. e113466.

37. Budden, K.F., et al., Emerging pathogenic links between microbiota and the gut-lung axis. Nat Rev Microbiol, 2017. 15(1): p. 55–63.

38. Brooks, B., et al., Microbes in the neonatal intensive care unit resemble those found in the gut of premature infants. Microbiome, 2014. 2(1): p. 1.

39. Quercia, S., et al., From lifetime to evolution: timescales of human gut microbiota adaptation. Front Microbiol, 2014. 5: p. 587.

40. Arrieta, M.C., et al., Early infancy microbial and metabolic alterations affect risk of childhood asthma. Sci Transl Med, 2015. 7(307): p. 307ra152.

41. Holt, P.G., The mechanism or mechanisms driving atopic asthma initiation: The infant respiratory microbiome moves to center stage. J Allergy Clin Immunol, 2015. 136(1): p. 15–22.

42. Stokholm, J., et al., Maturation of the gut microbiome and risk of asthma in childhood. Nat Commun, 2018. 9(1): p. 141.

43. Roussos, A., et al., Increased prevalence of irritable bowel syndrome in patients with bronchial asthma. Respir Med, 2003. 97(1): p. 75–9.

44. Rutten, N.B., et al., Intestinal microbiota composition after antibiotic treatment in early life: the INCA study. BMC Pediatr, 2015. 15: p. 204.

45. Vieira, W.A. and Pretorius, E. The impact of asthma on the gastrointestinal tract (GIT). J Asthma Allergy, 2010. 3: p. 123–30.

46. Wesemann, D.R. and Nagler, C.R. The Microbiome, Timing, and Barrier Function in the Context of Allergic Disease. Immunity, 2016. 44(4): p. 728–38.

47. Bassis, C.M., et al., Analysis of the upper respiratory tract microbiotas as the source of the lung and gastric microbiotas in healthy individuals. MBio, 2015. 6(2): p. e00037.

48. Man, W.H., de Steenhuijsen Piters, W.A., and Bogaert, D. The microbiota of the respiratory tract: gatekeeper to respiratory health. Nat Rev Microbiol, 2017. 15(5): p. 259–270.

49. Marsh, R.L., et al., The microbiota in bronchoalveolar lavage from young children with chronic lung disease includes taxa present in both the oropharynx and nasopharynx. Microbiome, 2016. 4(1): p. 37.

50. Burri, P.H., Fetal and postnatal development of the lung. Annu Rev Physiol, 1984. 46: p. 617–28.

51. German, R.Z. and Palmer, J.B., Anatomy and development of oral cavity and pharynx. GI Motil. Online http://www.nature.com/gimo/contents/pt1/full/gimo5.html, 2006.

52. Jimenez, E., et al., Staphylococcus epidermidis: a differential trait of the fecal microbiota of breast-fed infants. BMC Microbiol, 2008. 8: p. 143.

53. Hochberg, Z., et al., Child health, developmental plasticity, and epigenetic programming. Endocr Rev, 2011. 32(2): p. 159–224.

54. Manco, M., Gut microbiota and developmental programming of the brain: from evidence in behavioral endophenotypes to novel perspective in obesity. Front Cell Infect Microbiol, 2012. 2: p. 109.

55. Putignani, L., et al., The human gut microbiota: a dynamic interplay with the host from birth to senescence settled during childhood. Pediatr Res, 2014. 76(1): p. 2–10.

56. Ibberson, C.B., et al., Co-infecting microorganisms dramatically alter pathogen gene essentiality during polymicrobial infection. Nat Microbiol, 2017. 2: p. 17079.

57. Nakatsuji, T., et al., Antimicrobials from human skin commensal bacteria protect against Staphylococcus aureus and are deficient in atopic dermatitis. Sci Transl Med, 2017. 9(378).

58. Ramsey, M.M., et al., Staphylococcus aureus Shifts toward Commensalism in Response to Corynebacterium Species. Front Microbiol, 2016. 7: p. 1230.

59. Elhenawy, W., Debelyy, M.O., and Feldman, M.F., Preferential packing of acidic glycosidases and proteases into Bacteroides outer membrane vesicles. MBio, 2014. 5(2): p. e00909–14.

60. Koropatkin, N.M., Cameron, E.A., and Martens, E.C., How glycan metabolism shapes the human gut microbiota. Nat Rev Microbiol, 2012. 10(5): p. 323–35.

61. Rakoff-Nahoum, S., Coyne, M.J., and Comstock, L.E., An ecological network of polysaccharide utilization among human intestinal symbionts. Curr Biol, 2014. 24(1): p. 40–9.

62. Rakoff-Nahoum, S., Foster, K.R., and Comstock, L.E., The evolution of cooperation within the gut microbiota. Nature, 2016. 533(7602): p. 255–9.

63. Kim, M., et al., Gut Microbial Metabolites Fuel Host Antibody Responses. Cell Host Microbe, 2016. 20(2): p. 202–14.

64. Clemente, J.C., et al., The impact of the gut microbiota on human health: an integrative view. Cell, 2012. 148(6): p. 1258–70.

65. Knights, D., et al., Rethinking "enterotypes". Cell Host Microbe, 2014. 16(4): p. 433–7.

66. Ding, T., and Schloss, P.D., Dynamics and associations of microbial community types across the human body. Nature, 2014. 509(7500): p. 357–60.

67. Fukami, T., and Nakajima, M., Community assembly: alternative stable states or alternative transient states? Ecol Lett, 2011. 14(10): p. 973–84.

68. Gajer, P., et al., Temporal dynamics of the human vaginal microbiota. Sci Transl Med, 2012. 4(132): p. 132ra52.

69. Gibson, T.E., et al., On the Origins and Control of Community Types in the Human Microbiome. PLoS Comput Biol, 2016. 12(2): p. e1004688.

70. Gonze, D., et al., Multi-stability and the origin of microbial community types. ISME J, 2017. 11(10): p. 2159–2166.

71. Koren, O., et al., A guide to enterotypes across the human body: meta-analysis of microbial community structures in human microbiome datasets. PLoS Comput Biol, 2013. 9(1): p. e1002863.

72. Shetty, S.A., et al., Intestinal microbiome landscaping: insight in community assemblage and implications for microbial modulation strategies. FEMS Microbiol Rev, 2017. 41(2): p. 182–199.

73. Trosvik, P., and de Muinck, E.J., Ecology of bacteria in the human gastrointestinal tract– identification of keystone and foundation taxa. Microbiome, 2015. 3: p. 44.

74. Fadrosh, D.W., et al., An improved dual-indexing approach for multiplexed 16S rRNA gene sequencing on the Illumina MiSeq platform. Microbiome, 2014. 2(1): p. 6.

75. Caporaso, J.G., et al., QIIME allows analysis of high-throughput community sequencing data. Nat Methods, 2010. 7(5): p. 335–6.

76. Edgar, R.C., et al., UCHIME improves sensitivity and speed of chimera detection. Bioinformatics, 2011. 27(16): p. 2194–200.

77. DeSantis, T.Z, et al., Greengenes, a chimera-checked 16S rRNA gene database and workbench compatible with ARB. Appl Environ Microbiol, 2006. 72(7): p. 5069–72.

78. McDonald, D., et al., An improved Greengenes taxonomy with explicit ranks for ecological and evolutionary analyses of bacteria and archaea. ISME J, 2012. 6(3): p. 610–8.

79. Wang, Q., et al., Naive Bayesian classifier for rapid assignment of rRNA sequences into the new bacterial taxonomy. Appl Environ Microbiol, 2007. 73(16): p. 5261–7.

80. Nakagawa, S., and Schielzeth, H., A general and simple method for obtaining R2 from generalized linear mixed-effects models. Methods Ecol Evol, 2013. 4: p. 133–142.

